# Genomic epidemiology of enteropathogenic *Escherichia coli* (EPEC) in southwestern Nigeria

**DOI:** 10.1101/2025.02.14.638250

**Authors:** Olabisi C. Akinlabi, Rotimi A. Dada, Ademola A. Olayinka, Ibukunoluwa O. Oginni-Falajiki, Oyeniyi S. Bejide, Pelumi D. Adewole, Nicholas R. Thomson, Aaron O. Aboderin, Iruka N. Okeke

## Abstract

**Background:** Enteropathogenic *Escherichia coli* (EPEC) are etiological agents of diarrhea. We studied the genetic diversity and virulence factors of EPEC in southwestern Nigeria, where this pathotype is rarely characterized.

**Methodology/ Principal findings:** EPEC isolates (n=96) recovered from recent southwestern Nigeria diarrhea case-control studies were whole genome-sequenced using Illumina technology. Genomes were assembled using SPAdes and quality was evaluated using QUAST. Virulencefinder, SeroTypefinder, and Resfinder were used to identify virulence genes, serotypes, and resistance genes, while multilocus sequence typing was done by STtyping. Single nucleotide polymorphisms (SNPs) were called out of whole genome alignment using SNP-sites and a phylogenetic tree was constructed using IQtree. Thirty-nine of the 96 (40.6%) EPEC isolates were from cases of diarrhea. Nine isolates from diarrhea patients and four from healthy controls were typical EPEC, harboring bundle-forming pilus (*bfp*) genes whilst the rest were atypical EPEC. Fifteen isolates were EPEC-EAEC hybrids. Atypical serotypes O71:H19 (16, 16.6%), O108:H21 (6, 6.3%), O157:H39 (5, 5.2%), and O165:H9 (4, 4.2%) were most prevalent; only 8(8.3%) isolates belonged to classical EPEC serovars. The largest clade comprised ST517 isolates, of O71:H19 and O165:H9 serovars harboring multiple siderophore and serine protease autotransporter genes. An O71:H19 subclade comprised isolates <10 SNPs apart, representing a likely outbreak involving 15 children, four presenting with diarrhea.

**Conclusion/ Significance:** Likely outbreaks, of typical O119:H6(ST28) and atypical O127:H29(ST7798) were additionally identified. EPEC circulating in southwestern Nigeria are diverse and differ substantially from well-characterized lineages seen previously elsewhere. EPEC carriage and outbreaks could be commonplace but are largely undetected, hence, unreported, and require genomic surveillance for identification.

## INTRODUCTION

Enteropathogenic *Escherichia coli* (EPEC) was one of the first two diarrheagenic *E. coli* subcategories described and the pathotype was originally defined in the context of diarrhea outbreaks among young children (1). EPEC has more or less been eliminated as a public health threat in Europe and North America (2) but continues to cause infections in Asian, South American and African settings (3–5). However, it is infrequently reported from Africa (4,6,7) largely because of the difficulty in delineating EPEC from commensals. Earlier studies delineated EPEC by serotyping (8–10). However, this method does not capture EPEC that belong to serotypes that rarely or never circulated in the Western hemisphere since EPEC- identifying serotyping sets focus on only these so-called classical, serovarieties. Serovarieties represent clades descended from a common ancestor that produce similar O (somatic) and H (flagella) antigens (EPEC typically lack K antigens) (8). The Classical EPEC majorly belong to O serogroups (O55, O86, O111, O119, O125, O126, O127, O128, and O142) (9,11,12). However, within these serogroups, only certain serotypes are deemed classical EPEC, for example, O55:H6, O86:H34, O111:H2, O119:H2, O119:H6, O125:H6, O126:H2, O127:H^-^, O127:H6, O128:H2, and O142:H6 (11). While O51:H ^-^, O114:H2, O127:H40, O142:H34 are more recently discovered serotypes of typical EPEC that are often considered ‘classical’ (10). Identifying EPEC by their ability to produce attaching and effacing lesions on cultured epithelial cells represents a reliable but time- and resource-intensive means to identify this pathogen and has commonly been employed in South and Central America (13). Attaching-and-effacing is produced by a collection of virulence factors located on the Locus for Enterocyte Effacement (LEE) pathogenicity island (14). So-called typical EPEC (tEPEC) also carry one of several versions of a large virulence plasmid that includes genes encoding bundle-forming pili and the plasmid-encoded regulator (15). EPEC isolates may also express other virulence factors including, but not limited to non-LEE effectors (Nle) and autotransporters such as EspC, EspI, EspP, Pet, Pic and Sat (16).

Given the genetic basis for EPEC virulence is well understood, molecular methods to identify LEE and bundle-forming pilus genes represent the most commonly used approaches for its identification. We recently reported that PCR methods developed in other parts produced false-positives and negatives compared to whole genome sequencing in Nigeria (17). Here, we analyze whole-genome sequence-identified isolates from four diarrheal studies. We report a predominance of atypical EPEC isolates, belonging to non-classical serotypes, and retrospective identification of two outbreaks, one of which involved at least 19 children.

## METHODOLOGY

### Ethical considerations

Ethical approval for this research was received from UI/UCH Ethics Committee with assigned number UI/EC/15/0093, UI/EC/18/0335 and from Obafemi Awolowo University Teaching Hospital Complex with assigned number IRB/IEC/0004553.

### Isolates analyzed

This study analyzed the genomes of 96 EPEC isolates isolated from four different case-control studies. In two of the studies, stool specimens were collected from children with diarrhea and healthy controls. Three of these studies were conducted in Ibadan, Oyo State [one unpublished and (6,7) and the fourth in Ile-Ife in adjoining Osun State (18). The inclusion criteria for three studies on childhood diarrhea have been published (6, 17). A fourth study, still in analysis, collected specimens from people living with HIV, largely adults and from HIV negative patients with diarrhea visiting two health care facilities in Ibadan.

### Whole genome Sequencing

EPEC isolates from children in Ile-Ife were whole genome sequenced using the same protocol employed for sequencing the Ibadan isolates (19). Briefly, *E. coli* genomic DNA was isolated using the Wizard Genomic Extraction kit (Promega) according to the manufacturer’s instructions, The DNA extracts were quantified using a dsDNA broad range quantification assay (Invitrogen) and transported to the Wellcome Trust Sanger Institute for sequencing on the Illumina platform. The SPAdes assembler was used to assemble raw data in Nigeria, and the assembly quality was assessed using the Quality Assessment Tool for Genome Assemblies (QUAST) and CheckM. Speciation was performed using BactInspector.

### Whole genome sequence analyses

Resistance genes were identified using the ARIBA and Resfinder databases (20). Virulence factors and plasmid replicons were detected using VirulenceFinder and Plasmidfinder, respectively (21,22). Isolates carrying any *eae* allele but lacking Shiga toxin genes, were classed as EPEC. Intimin alleles were delineated using BLAST (23). Multilocus sequence typing (MLST) was performed using STtyping (24). *In silico* serotyping was carried out using Ectyper (25). Isolates belonging to serogroups, O55:H6, O86:H34, O111:H2, O119:H2, O119:H6, O125:H6, O126:H2, O127:H^-^, O127:H6, O128:H2, O142:H6 (11)O51:H ^-^, O114:H2, O127:H40, O142:H34 were classed as classical EPEC and others, non-classical (11,26). Sequence data were submitted to ENA and are available as Bioproject PRJEB8667 at ENA https://www.ebi.ac.uk/ena/browser/home and Genbank https://www.ncbi.nlm.nih.gov/genbank/.

### Nutritional status

Nutritional status at the time isolates were recovered was determined by computing weight for height, height for weight and weight for age from data collected with the specimens and interpreting them according to WHO criteria https://www.who.int/data/nutrition/nlis/info/malnutrition-in-children

## RESULTS

### Classification of EPEC isolates

A total of 96 EPEC isolates were isolated from all four studies, 39 from individuals with diarrhea and 57 from healthy controls. EPEC were isolated from children presenting at Ibadan’s University Teaching Hospital Ibadan (UCH) - 9 (9.4%), Primary Health Care centers in Ibadan 24 (25.0%), and primary, secondary, and tertiary hospitals 25(26.0%) in Ile-Ife (Table 1). Adult samples were from PLWHIV attending UCH’s Infectious Disease Institute 3(3.1%) or HIV-negative patient controls from other UCH clinics, as shown in Table 1. Eleven of the isolates were from children under one year of age with diarrhea and 19 from healthy children in the same age bracket. EPEC were not associated with diarrhea among study participants in any age range, including children under 1 year, children aged 0-6 months and children aged 7-12 months (Table 1)

**Table 1:** Age distribution of individuals from whom EPEC were recovered.

|  | Cases (n=975) | Controls (n=932) | Total (n=1907) | p-value |
| --- | --- | --- | --- | --- |
| 0 to 3 months | 6 | 4 | 10 | 0.806 |
| 4 to 6 months | 0 | 5 | 5 | 0.0654 |
| 7 to 9 months | 3 | 6 | 9 | 0.4616 |
| 10 to 12 months | 2 | 4 | 6 | 0.6424 |
| 13 and above | 8 | 16 | 24 | 0.0998 |
| Adult | 3 | 0 | 3 | 1.0000 |

More - 57 (59.4%) - of the EPEC isolates were from healthy individuals (Table 2) and the majority 83(3.9%) of isolates were atypical EPEC, belonging to non-classical EPEC serotypes. However, typical EPEC, which were uncommon overall, were recovered from 9 (1.1%) individuals with diarrhea and 4 (0.3%) apparently healthy controls (p<0.05, Table 3).

**Table 2:** EPEC isolates isolated from recent southwest Nigeria epidemiological studies.

| Source studies | Study Type | Cases with diarrhea | Controls without diarrhea | Total | Cases | Controls | EPEC n (%) | p-value |
| --- | --- | --- | --- | --- | --- | --- | --- | --- |
| Ibadan Primary Health Care Center (PHC) (27) | Children aged under 5 | 120 | 357 | 477 | 7 | 17 | 24(5.0%) | 0.6331 |
| University Teaching Hospital Ibadan (UCH) (unpublished) | Children aged under 5 | 359 | 241 | 600 | 6 | 3 | 9(1.5) | 0.9372 |
| Ife primary, secondary, and tertiary hospitals (IFE) (18) | Children aged under 5 | 167 | 334 | 501 | 8 | 17 | 25(5.0%) | 1.000 |
| Infectious Disease Institute (IDI) (7) and unpublished | People living with HIV | 202 | N/A | 202 | 1 | N/A | 3(0.9%) | 0.6837 |
| Infectious Disease<br>Institute (IDI) (7)<br>and unpublished | HIV-negative<br>individuals | 127 | N/A | 127 | 2 | N/A |  |  |
| Total |  | 975 | 932 | 1907 | 24 | 37 | 61(3.2%) |  |

**Table 3:** Identified EPEC from the studies.

|  | Case isolates<br>from patients<br>N=805<br>(%) | Control isolates from<br>symptomless individuals<br>N=1311<br>(%) | Total isolates<br>from 2116<br>individuals<br>(%) | p-value |
| --- | --- | --- | --- | --- |
| All EPEC | 39 (4.8) | 57 (4.3) | 96(4.5) | 0.6703 |
| Typical | 9 (1.1) | 4 (0.3) | 13(0.6) | **0.0417** |
| Atypical | 30 (3.7) | 53 (4.0) | 83(3.9) | 0.8040 |
| Classical | 5 (0.6) | 3 (0.2) | 8(0.4) | 0.2879 |
| Non-classical | 34(4.2) | 54(4.1) | 88(4.2) | 0.9111 |

### Phylogroup, ST, intimin type and Serogroups of the EPEC isolates

Forty-six (47.9%) of the EPEC isolates belonged to *E. coli* phylogroup B1. Of the others 20 (20.8%), 17 (17.7%), 5 (5.2%), 1 (1%) were from phylogroups B2, A, D and E, respectively. In addition, 3 (3.1%) were untypeable and none belonged to phylogroups C or F (Table 3). All the identified typical EPEC belonged to phylogroups A, B1, and B2 none from D, E, and U. while all classical EPECs belonged to phylogroup B2 (Figure 1).

**Fig 1:**
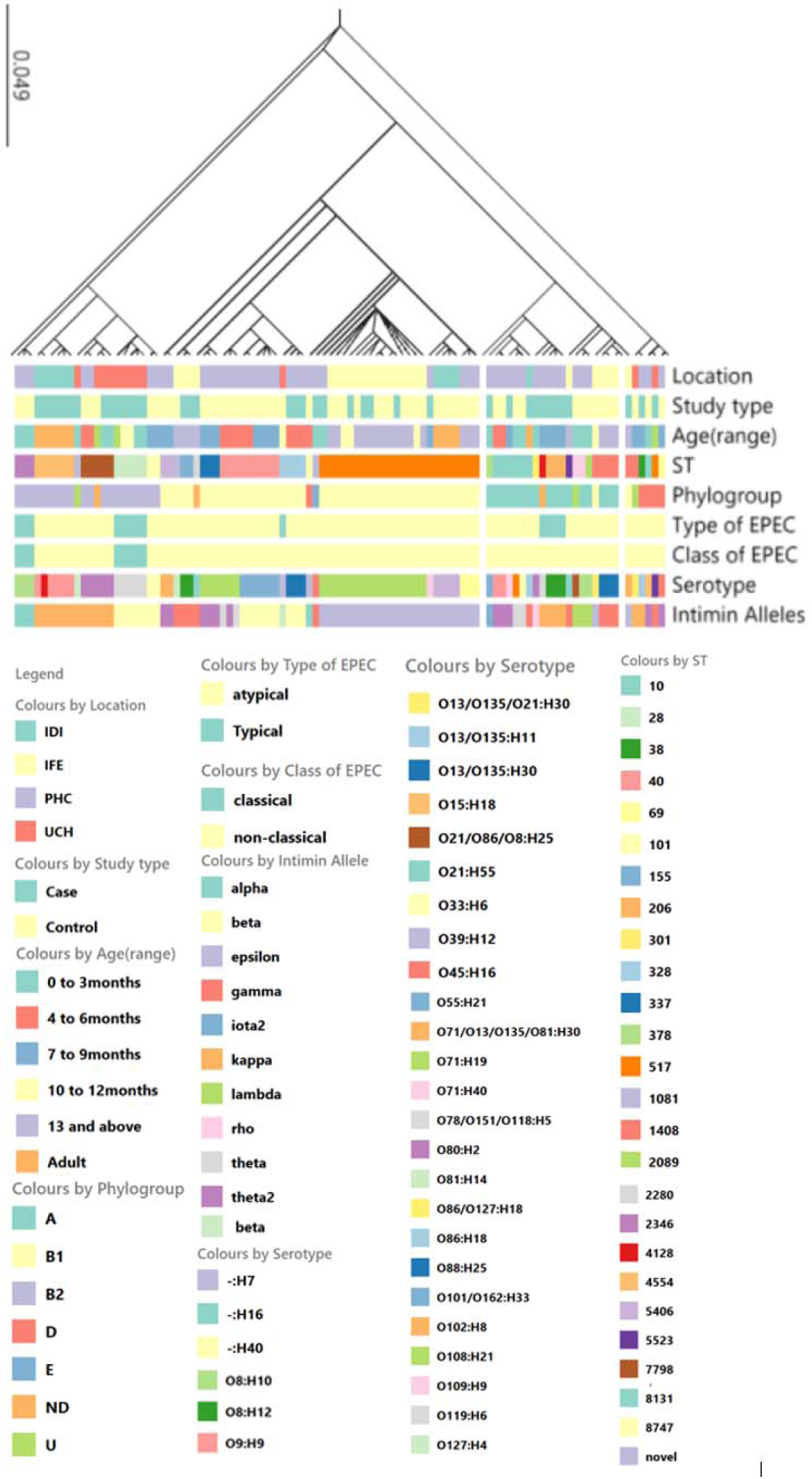
Maximum-likelihood whole-genome SNP-based tree of EPEC isolates from this study juxstaposed against source (location, age and health status of patient) and classification (typical/ atypical, classical or non-classical serovar, intimin type, phylogroup and ST) criteria. Data are available in interactable format at https://microreact.org/project/hnPuLKV1d7pnskcG146MJj-epec-isolates-and-their-locations-2025

The EPEC isolates belonged to 26 different STs with ST 517 being the most common. ST517 isolates were recovered from three different recruiting sites (IFE, UCH and PHC) as shown in Figure 1. Typical EPEC isolates belonged to ST28, ST206 (and related ST4128) or ST7798. Atypical EPEC were distributed among the seven STs illustrated in Figure 1. Classical EPEC belonged to two STs: ST28 (O119:H6) and ST2346 (O142:H34), all ST28 EPEC were from children with diarrhea while ST2346 were all from healthy children. The classical isolates were all typical EPEC carrying both the *bfp* and *eae* gene (Figure 1).

A total of 39 serotypes were identified with serotypes O71:H19 as the most prevalent 16 (16.7%). EPEC belonging to O71:H19 serotype were isolated from Ile-Ife (IFE) and Ibadan Primary Health Centers (PHC) (Figure 1). The typical EPEC isolates belonged to classical serotypes O119:H6 (n=53), O142:H34 (n=3) and non-classical O151/O118:H5 (n=4) were all typical EPEC while other serotypes were atypical EPEC.

Based on genome data ten different intimin types were identified among the EPEC isolates with epsilon being the most prevalent 26 (27.1%), following beta 17(17.7%), theta2 and gamma 11(11.4%) (Table 6). All typical EPEC isolates carried intimin beta or kappa but a much broader range of intimin alleles was seen among atypical isolates (Figure 1).

None of the isolates in the study belonged to the earliest EPEC phylogenomic groups, originally delineated by multilocus enzyme electrophoresis, EPEC1 and EPEC2. MLST and then whole genome sequence approaches have since confirmed these EPEC lineages and uncovered a total of 15 EPEC phylogenomic groups (28). Most of the isolates in this study belong to new phylogenomic lineages that are yet to be described in the literature, however, some belonged to EPEC4 - 7 (7.7%), EPEC5 - 3 (3.3%), EPEC7 - 4 (4.4%), EPEC9 - 3 (3.3%), and EPEC10 – 7 (7.7%). The majority of our isolates are closely related to isolates from Gambia belonging to an as yet unnamed lineage (28) (Figure 2).

**Fig 2:**
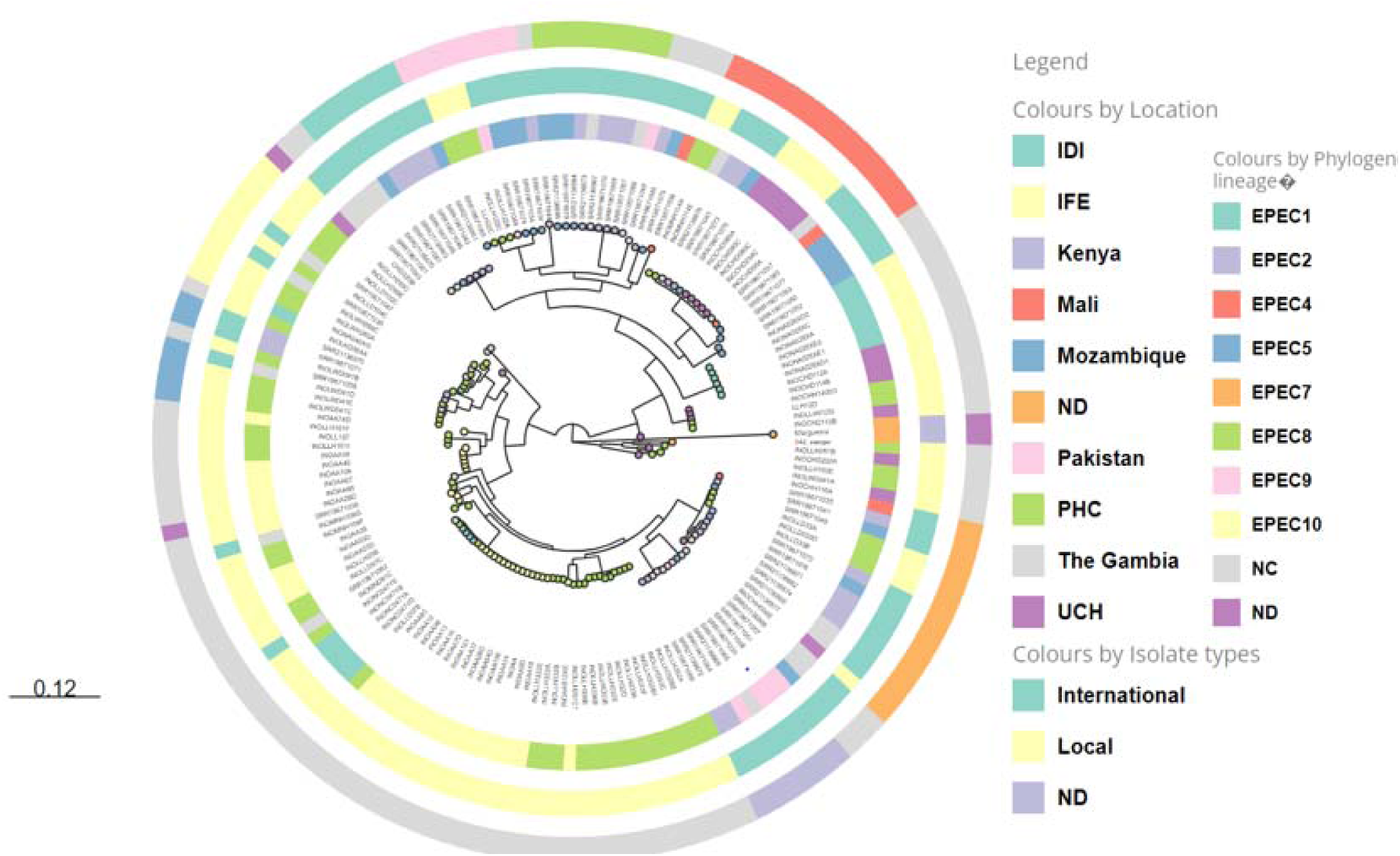
Maximum-likelihood whole-genome SNP-based tree and results of phylogenomic analysis of 61 EPEC isolates from Nigeria in this study and 67 international strains. The inner circle represents the isolation sources, the middle circle represents the strain types, with typical in green, outer circle represent the phylogenomic lineages, ND; not determined Local isolates, NC; not classified international strains. Data are available in interactable format at: https://microreact.org/project/usSgwCwVrm1AZ76kHqfk1d-epec-local-and-international-strains

### EPEC Hybrids

To determine if the EPEC identified here carried genes associated with other *E. coli* pathotypes we searched their genomes using Virulencefinder. None of the isolated EPEC isolates carried Shiga toxin genes – no enterohaemorrhagic *E. coli* were recovered in any of the epidemiological surveys. However, most of the studies yielding the isolates were focused on infants, from whom EHEC are rarely recovered. However, one O101/O162:H33 isolate carried the enterohemolysin gene, *ehx*, which is typically contained on an EHEC virulence plasmid. This ST378 phylogroup isolate is the only one bearing the Iota allele of intimin and is related to a recently described atypical EPEC10 clone from Brazil shown to induce mucus hypersecretion (29). The isolate was recovered from a child attending an Ibadan primary health center. Isolates belonging to this clone carry aerobactin and yersiniabactin siderophore genes, a range of non-LEE effector (*nle*) genes and *hlyE* but lack SPATE genes (29). Our own O101/O162H:33 isolate fit this profile and also carried enteroaggregative *E. coli* (EAEC) aggregative adherence fimbriae I (*agg*) genes and their regulator *aggR*. Thus, it is a hybrid EPEC-EAEC isolate, although it lacked other virulence genes common in EAEC such as SPATEs, the antiaggregative protein (*aap*) gene and its secretion system encoded by *aat* genes (Figure 3).

**Fig 3:**
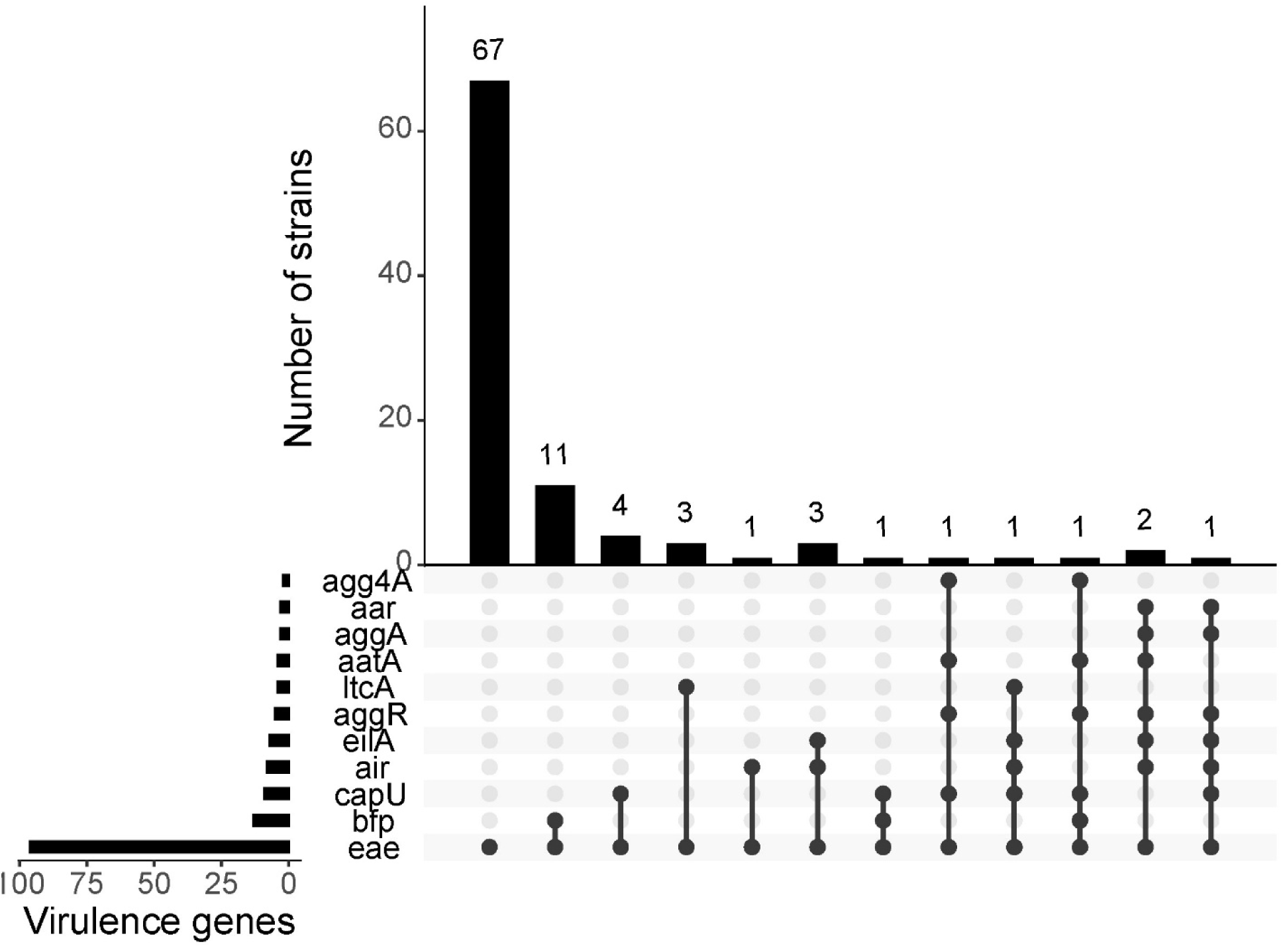
EPEC hybrids carrying EAEC (*air, eilA, aggR, aap, aggA, agg4A, aar* and *capU*) and/or ETEC (*ltcA*) virulence genes

A total of 15 EPEC isolates evaluated in this study, including the O101/162:H33 isolate, co-harbored EAEC virulence genes (Figure 3) such as the EAEC hexosyltransferase homolog *(capU)* - 9(8%), enteroaggregative immunoglobulin repeat protein (*air) -* 8(8%)*, Salmonella* HilA homolog (*eilA*) - 7(7%), aggregative adherence regulator *(aggR) -* 5(5%), anti-aggregation protein transporter (*aatA) -* 4(4%), AAF/I fimbrial subunit *(aggA) -* 3(3%) and AAF/IV fimbrial subunit (*agg4a*) *-* 2(2%). Genes encoding the commonly encountered AAF/II, AAF/III and AAF/V fimbriae were not found in any hybrid isolate.

Four of the isolates represented EPEC- enterotoxigenic *E. coli* (ETEC) hybrids, carrying heat labile endotoxin virulence genes (*ltcA*) were identified and one of these was an EPEC-EAEC- ETEC hybrid that also carried *eilA*, air and *capU* genes (Figure 1). Five isolates carried *afa* genes. The Afa/Dr fimbriae define diffusively adherent *E. coli* (DAEC) (but are also found in other pathotypes) and are associated with IL-8 secretion in diarrhea, a marker of inflammation. Three of these isolates were EPEC-EAEC hybrids and one of them was the EPEC-EAEC-ETEC hybrid (Figure 3).

Altogether, 18 (18.8%) isolates in the study carried, in addition to the EPEC virulence genes, genes associated with other diarrheagenic *E. coli* pathotypes. Only one was a typical EPEC isolates, positive for EAF plasmid genes *bfp* and *per*. Hybrid genotypes were distributed across the phylogeny (five, five, five, one and two, respectively belonged to *E. coli* phylogroups A, B1, D, E and untypable). Nine of the eighteen hybrid isolates were recovered from children with diarrhea.

### EPEC outbreaks

Multiple EPEC clades comprised genomes separated with 20 or less SNPs. ST28 EPEC isolates (O119:H6) were recovered from three pediatric patients presenting at the same referral hospital. A cluster of five atypical EPEC serogroup O127:H29 ST7798 isolates were from patients, all aged 3 months, attending primary health care centers (n=2) and the referral hospital (n=3) in Ibadan. Two of the children at the referral center presented with diarrhea in 2017 and one was a healthy control. These isolates differed by only 2 SNPs and the time and place data suggest that they are epidemiologically connected and so represent a localized outbreak. Interestingly two apparently healthy children recruited from an Ibadan primary health care centre in 2015 yielded very similar ST7798 isolates, being only 10-11 SNPs away from the hospital isolates suggesting that this ST7798 atypical lineage may be circulating more broadly in the community across Ibadan and that confirming EPEC outbreaks with genomic information alone may not be possible.

ST 517 EPEC was the most prevalent (n = 25, 26%) ST from this study and they were all atypical (Figure 1). As shown in Figure 2, ST 517 EPEC was isolated from both cases and control children’s specimens from IFE, PHC and HIV patients in IDI but none from UCH. All ST517 isolates carried intimin epsilon genes. There are three sub-clades within the ST517 clade. The first is comprised of O171:H19 isolates from healthy children attending primary health centers in Ibadan and is related to the second clade of O165:H9 and O109:H9 isolates from children or PLWHIV attending clinics in Ibadan. The third and largest clade is comprised of O171:H19 isolates from children recruited in Ife. Four of these were from children with diarrhea and 11 were from controls (Figure 4). When the number of SNP distances among these isolates was computed, it was observed that the ST517 EPEC clade is made up of isolates <10 SNPs apart and they were all recovered from specimens obtained between the 4^th^ of August and the 21^st^ of September 2017 (Figure 4a). Three of the four case isolates and six of the eleven isolates from asymptomatic individuals were all recovered from specimens collected on the 28^th^ of August 2017 and all of them resided within a circle with a 1.2 km radius.

**Fig 4:**
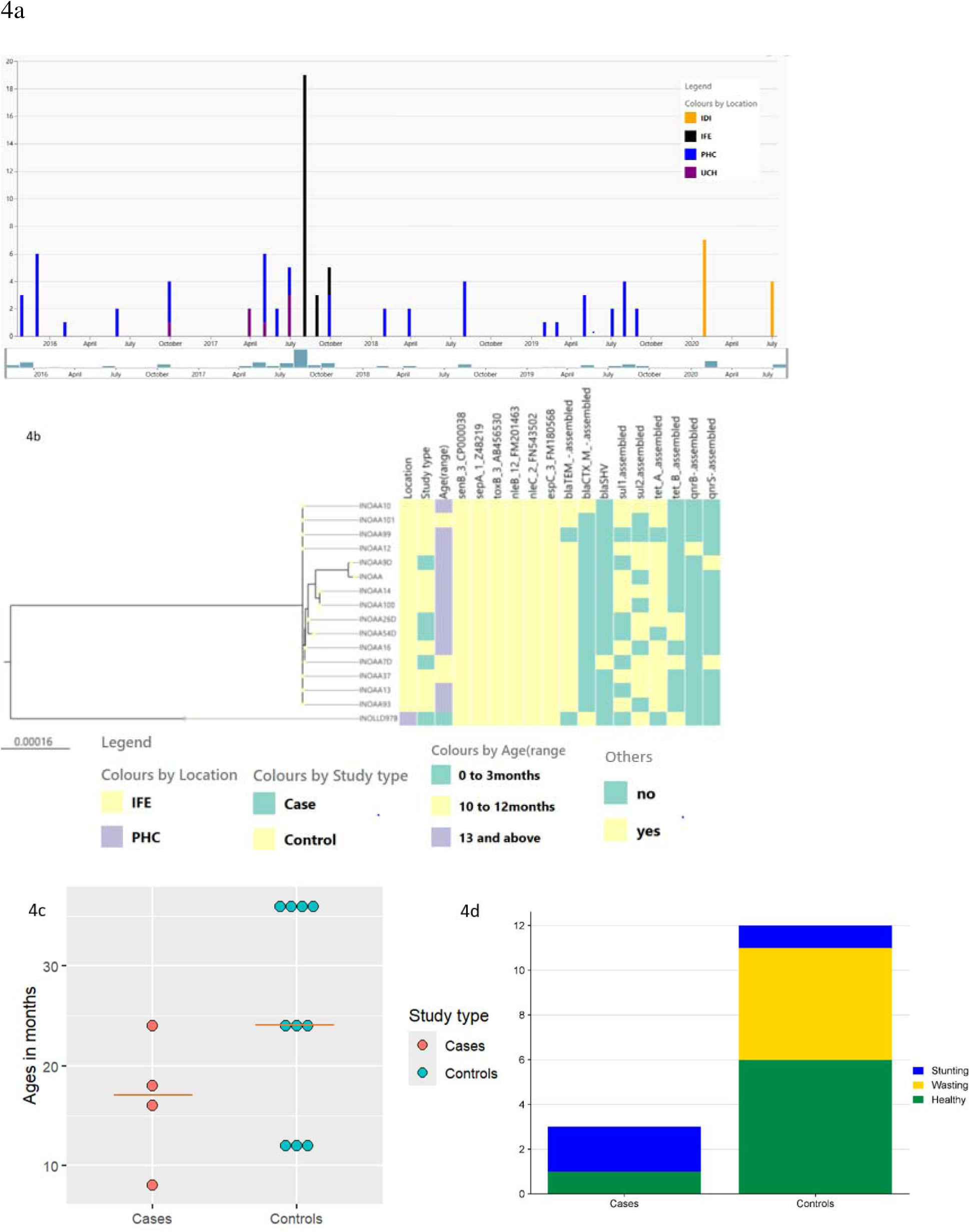
(A). Epicurve showing the dates of enrollment of children with diarrhea (top) and those from whom ST517 isolates were recovered (bottom). (B). Virulence and antimicrobial resistance genes in ST517 isolates from Ife that, based on sharing <10 SNPs, are believed to constitute an outbreak. (C). Age distribution of individuals from whom EPEC included in the tree in (B) were recovered with the red line representing the median age for each category. (D). The nutritional status of children from whom isolates in (B) above were collected.

All the ST517 likely outbreak isolates from Ile-Ife had, in addition to a LEE carrying intimin epsilon, the *sepA, senB, toxB, espC, nleB* and *nleC* virulence genes. They also harbored antimicrobial resistance genes conferring resistance to extended-spectrum beta-lactam drugs, sulfonamide and tetracycline (Figure 4b). Notably, no virulence loci were more predominant in case isolates, compared to controls (p > 0.05).

The children from the likely outbreak who had diarrhea were aged 8, 16, 18 and 24 months. Three of the children without diarrhea were aged 12 months and the rest were aged 24-36 months (Figure 4c). While undernutrition was common as a whole proportionately more of the children who were not stunted carried ST517 EPEC outbreak isolates without symptoms. Two children who succumbed to diarrhea showed significantly lower weights for age and heights for age than healthy children stunted growth (Figure 4d).

## DISCUSSION

Diarrheagenic *E. coli* are prominent causes of diarrhea in children under five years of age with ETEC, EAEC, and EPEC isolates being the most common (30). EPEC was among the first pathotypes delineated, has been eliminated as a significant cause of infantile diarrhea in North America and Europe, but remains common in Africa (31–35). Sequence-based characterization of EPEC from Africa is uncommon and therefore little is known about circulating lineages, particularly after classical group serotyping, which was notably insensitive in Africa, was no longer advocated for routine use (36). In this study, which identified and subtyped EPEC by whole genome sequencing, typical EPEC, and EPEC belonging to classical serotypes, were uncommon. Majority of the EPEC isolates examined in this study 67 (69.8%) belong to newly identified phylogenomic lineages, in keeping with reports of EPEC from recent Africa and Asia studies (5,37,38). Our findings highlight the diversity of the EPEC pathotype, reflect its convergent origins and continuous evolution, and demonstrate that more studies need to be conducted in the parts of the world most burdened by EPEC disease (5,37–39).

Enteropathogenic *E. coli* are classified as typical or atypical EPEC based on the presence of the EPEC adherence factor (EAF) plasmid, which carries *bfp* and *per* genes (26). The diarrheagenicity of typical EPEC isolates has been confirmed in adult volunteer challenge studies (40), but less evidence is available to understand the virulence of atypical strains (41). However, atypical strains belong to far more lineages and carry a broad range of potential virulence loci, so more research is likely to delineate additional hypervirulent lineages and virulence genes. In this study, as in many other recent African studies (42), the EPEC isolates detected from Ibadan and Ife studies were mostly atypical EPEC 83(86.5%), which are emerging enteropathogens with global distribution (43–45). They may be replacing typical EPEC as a leading cause of diarrhea in both developing and industrialized nations (43–45). Their historical epidemiology, particularly in Africa, is difficult to understand since earlier studies used diagnostic methods biased towards typical EPEC. Typical EPEC were uncommonly recovered in this study, but were more commonly isolated from children with diarrhea (p<0.05) and four of the seven affected children were enrolled at referral centres, suggesting disease severity.

EPEC were earlier classified based on their somatic (O) and flagellar (H) antigens which have been associated with diarrhea (46). The World Health Organization designated 12 EPEC serotypes in 1987: O26, O55, O86, O111, O114, O119, O125, O126, O127, O128, O142, and O158 (26). Strains belonging to some of these serogroups are known as classical EPEC. More classical EPEC are typical but strains belonging to these serogroups include atypical EPEC isolates and other pathotypes of DEC, such as EAEC (26,47,48). Specific O:H types are more reliable markers of EPEC and often represent lineages disseminated globally (8,11,12). In this study, we recovered O119:H6 and O142:H34 isolates, which represent well-described serotypes of typical EPEC (10,49,50) as well as two EPEC-EAEC hybrids belonging to O86:H18 a serovar originally described as EPEC and then re-classified as EAEC (51). A number of isolates belonged to O-groups commonly associated with EHEC, including O55, O157 and O118 were isolated but none of these isolates had Shiga-toxin genes, adding to the evidence that sub-classifying DEC by serotyping in our setting could lead to pathotype miscalls (52,53). Majority, 88(91.7%), of the identified EPEC in this study were non-classical EPEC.

It is challenging to find historical context of the data gathered in our study because previous studies in Nigeria lack whole genome-sequence data for EPEC or indeed archived the isolates. Serotyping is the most commonly employed delineation tool. Unless performed at a reference centre, it can be inaccurate and only finds strains belonging to classical groups or types. (31) We can glean some learnings when we juxtapose our data on data from studies that used robust serotyping methods. Ifeanyi et al. (2017) identified many EPEC serovarieties and recovered many isolates they deemed non-typeable, suggested that they might belong to non-familiar serotypes. While rarely reported elsewhere, O33:H6/H34 EPEC were found at low frequency in all studies (31, 32), including this one (belonging to ST8747), indicating the possibility that they represent a local endemic clade.

The intimin gene, encoded by the *eae* gene that is part of the LEE, is required for intestinal colonization in the host and the EPEC-defining attaching and effacement phenotype. Different intimin subtypes are classified based on the variable 280-amino acid C-terminal sequence of intimin (Int_280_) (54,55). The most prevalent intimin subtypes among aEPEC isolates of various serotypes worldwide have been reported to be Alpha (α), Beta (β), Gamma (γ), Zeta (ζ), Delta (δ) and Epsilon (□), while other intimin subtypes are said to be less common (34,55,56). In this study all the stated intimin types were identified except for Delta and Zeta. Moreover, 38(39.6%) of the isolates carried Iota2, Kappa, Lambda, Rho, Theta and Theta2 subtypes. The occurrence of different intimin subtypes confirms the diversity of EPEC isolates circulating in Nigeria and their under-representation among well-studied strains. We have previously reported that consensus intimin PCR primers prime Iota, Rho, and Theta alleles with low efficiency, and their abundance in our strain set emphasizes the need to develop diagnostic tools that are inclusive of or specific for prominent West African lineages. We note that the intimin Iota2-bearing ST378 isolate we identified is closely related to a hypervirulent clone reported from Gambia (4) and therefore these lineages unfamiliar to science may have more than local significance.

The number and variety of hybrid isolates detected in this study demonstrates the complex evolutionary dynamics of EPEC, and the *E. coli* species more generally. Hybrid EPEC were more common than, and did include, EPEC isolates carrying the EPEC adherence plasmid that defined typical EPEC. In this regard, EPEC carrying virulence genes other than *bfp* have greater epidemiological significance in our setting. They were found throughout the phylogeny, evolution by independent horizontal acquisition of relevant virulence genes may be common. Nine of 18 hybrid isolates were from children with diarrhea, and therefore hybrids (and how they evolve) should be the subject of future research.

We identified three clonal clusters: ST28 (O119:H6, typical EPEC) and an ST7798 (O127:H29, atypical EPEC) appear to be connected to an outbreak, both including children presenting at a referral hospital in Ibadan, as well as an ST517 outbreak that was picked up from children attending an Ile-Ife primary health care center. All three likely outbreaks revealed that healthy carriers as well as ill children were colonized. The most prominent lineage in this study was an ST517 EPEC clade of closely related isolates that was shown to represent the largest outbreak that occurred in Ile-Ife in August/September 2017. ST517 strains are not commonly reported in the literature and this clade has yet to be assigned an EPEC phylogroup. One report, from China, implicated imported foods (57) and O71:H19 EPEC has been identified from human stool samples in Brazil, South Korea and in dog feces from Minnesota (58–60), implying that O71:H19/ ST571 EPEC are broadly disseminated and may colonize and/or cause disease humans and animals. As many as 17 children cultured in that study carried the strain but only four were diarrhea patients (18). It was observed that there were no easily apparent differences among the bacterial isolates from children with diarrhea and asymptomatic children in this study. However, those children with diarrhea were stunted in growth. Altogether, our data suggest that EPEC outbreaks can occur undetected and that when they do, the most vulnerable are at highest risk.

## CONCLUSION

EPEC isolates circulating in Ile-Ife and Ibadan, Nigeria are diverse and include globally disseminated clones of known virulence as well as EPEC that differ substantially from well-characterized lineages reported from other parts of the world. EPEC isolates that carry EAEC, ETEC or DAEC genes are common in our study and the current definitions that classify all EAF/*bfp*-negative strains as ‘atypical’ support the idea that very little is known about the pathogenesis of EPEC lineages, which are endemic in southwestern Nigeria. Genomic surveillance can uncover EPEC outbreaks, the scale and frequency of which are largely unknown and grossly under-reported. As infected people may be asymptomatic and our data suggest that vulnerable individuals such as nutritionally deficient children may bear the brunt of EPEC disease, there is need for active, continued genomic surveillance of the epidemiology of EPEC in our setting.

## Acknowledgement

We thank Stella Ekpo, Amos Olowookere, Justice C Onwuka, Mariam Odebode, Elizabeth Akande and El-shama Q Nwoko for technical assistance. We are grateful to clinicians, notably Drs A Adepoju, BO Ogunbosi, KO Akande and T Ilori, and consenting patients who enabled the epidemiological studies from which the isolates were obtained.

